# New Insights into the Mechanisms Used by Inhibitors Targeting Glutamine Metabolism in Cancer Cells

**DOI:** 10.1101/2021.09.20.461106

**Authors:** Shawn K. Milano, Qingqiu Huang, Thuy-Tien T. Nguyen, Sekar Ramachandran, Aaron Finke, Irina Kriksunov, David Schuller, Marian Szebenyi, Elke Arenholz, Lee A. McDermott, N. Sukumar, Richard A. Cerione, William P. Katt

## Abstract

Many cancer cells become dependent on glutamine metabolism to compensate for glycolysis being uncoupled from the TCA cycle. The mitochondrial enzyme Glutaminase C (GAC) satisfies this ‘glutamine addiction’ by catalyzing the first step in glutamine metabolism, making it an attractive drug target. Despite one of the allosteric inhibitors (CB-839) being in clinical trials, none of the drugs targeting GAC are approved for cancer treatment and their mechanism of action is not well understood. A major challenge has been the rational design of better drug candidates: standard cryo-cooled X-ray crystal structures of GAC bound to CB-839 and its analogs fail to explain their potency differences. Here, we address this problem by using an emerging technique, serial room temperature crystallography, which enabled us to observe clear differences between the binding conformations of inhibitors with significantly different potencies. A computational model was developed to further elucidate the molecular basis of inhibitor potency. We then corroborated the results from our modeling efforts by using recently established fluorescence assays that directly read-out inhibitor binding to GAC. Together, these findings provide new insights into the mechanisms used by a major class of allosteric GAC inhibitors and for the future rational design of more potent drug candidates.

## Introduction

One of the most recognized phenotypes of many cancer cells is a metabolic shift from oxidative phosphorylation to aerobic glycolysis, commonly described as the Warburg effect (Vander Heiden, Cantley, & Thompson, 2009). Cells undergoing aerobic (Warburg) glycolysis make use of additional sources of carbon, such as glutamine, which exists in high concentrations in blood plasma (Lukey, Katt, & Cerione, 2017). Cancer cells often overexpress glutaminase enzymes, in particular glutaminase C (GAC), which resides in the mitochondria and catalyzes the hydrolysis of glutamine to glutamate. Glutamate is then either used to fuel the TCA cycle via its conversion to α-ketoglutarate by glutamate dehydrogenase or as a building block for various biomolecules (William P Katt, Lukey, & Cerione, 2017). High levels of GAC have been observed in aggressive cancers and the inhibition of its enzymatic activity has been shown to reduce the proliferative capability of a variety of different cancer cells, and often their survival, both *in vitro* and in mouse models (M. I. Gross et al., 2014; W.P. Katt, Ramachandran, Erickson, & Cerione, 2012; Wang et al., 2010). Moreover, GAC inhibitors have been shown to improve sensitivity to different clinical drug candidates, including the recent demonstration that their combination with antibodies targeting the immune checkpoint protein PD-L1 offers exciting therapeutic potential (Byun et al., 2020; M. Gross et al., 2016; Varghese et al., 2021). These findings have led to sustained interest in examining GAC as an anti-cancer drug target, and that inhibitors targeting this enzyme may find clinical relevance after suitable development.

Numerous GAC inhibitors have been reported, with the most heavily investigated being a class of compounds derived from the small molecule BPTES (bis-2-(5-phenylacetamido-1,3,4-thiadiazol-2-yl)ethyl sulfide) (Hartwick & Curthoys, 2011; Robinson et al., 2007). These compounds bind at the dimer/dimer interface of the GAC tetramer, which is near the so-called activation loop (Gly315 to Glu325), effectively trapping the tetrameric enzyme in an inactive conformation (DeLaBarre et al., 2011; Ferreira et al., 2013; Hartwick & Curthoys, 2011; Qingqiu Huang et al., 2018; Li et al., 2019; Robinson et al., 2007; Stalnecker et al., 2014; Thangavelu et al., 2012). Over 2,000 BPTES analogs have been reported to date, primarily in the patent literature, with BPTES and CB-839 (Fig. 1A) being the most studied (M. I. Gross et al., 2014; William P Katt et al., 2017). Both CB-839 and a more recently described compound, IPN60090, have advanced to clinical trials (Harding et al., 2015; Soth et al., 2020), although CB-839 has not been approved as yet as an anti-cancer drug, while IPN60090 has reportedly been removed from trials entirely (AdisInsight, 2021). Despite the extensive optimization efforts conducted during its discovery, CB-839 has both a higher calculated logP (ClogP, a measure of lipophilicity) and lower lipophilic efficiency (LipE, a measure of inhibitory potency relative to lipophilicity) than BPTES (L. A. McDermott et al., 2016). We developed the UPGL series of inhibitors with the aim of replacing the flexible linker present in BPTES and CB-839 with a rigid, saturated heterocyclic ring as a means of improving the physicochemical properties of the drugs by minimizing the number of rotatable bonds and increasing potency via the reduction of the entropic penalty to protein binding. Some of these compounds have been able to surpass CB-839 in potency, LipE, and resistance to degradation by liver microsomes (L. A. McDermott et al., 2016). However, the X-ray crystal structures that we previously determined for the five potent UPGL molecules shown in Fig. 1B, bound to GAC (Qingqiu Huang et al., 2018; L. A. McDermott et al., 2016), as well as those reported for the enzyme complexed either to CB-839 or to the less potent compound BPTES (DeLaBarre et al., 2011; Qingqiu Huang et al., 2018; Thangavelu et al., 2012), are all very similar and thus provide little insight into the mechanistic basis of inhibition and what regulates potency.

**Figure 1:**
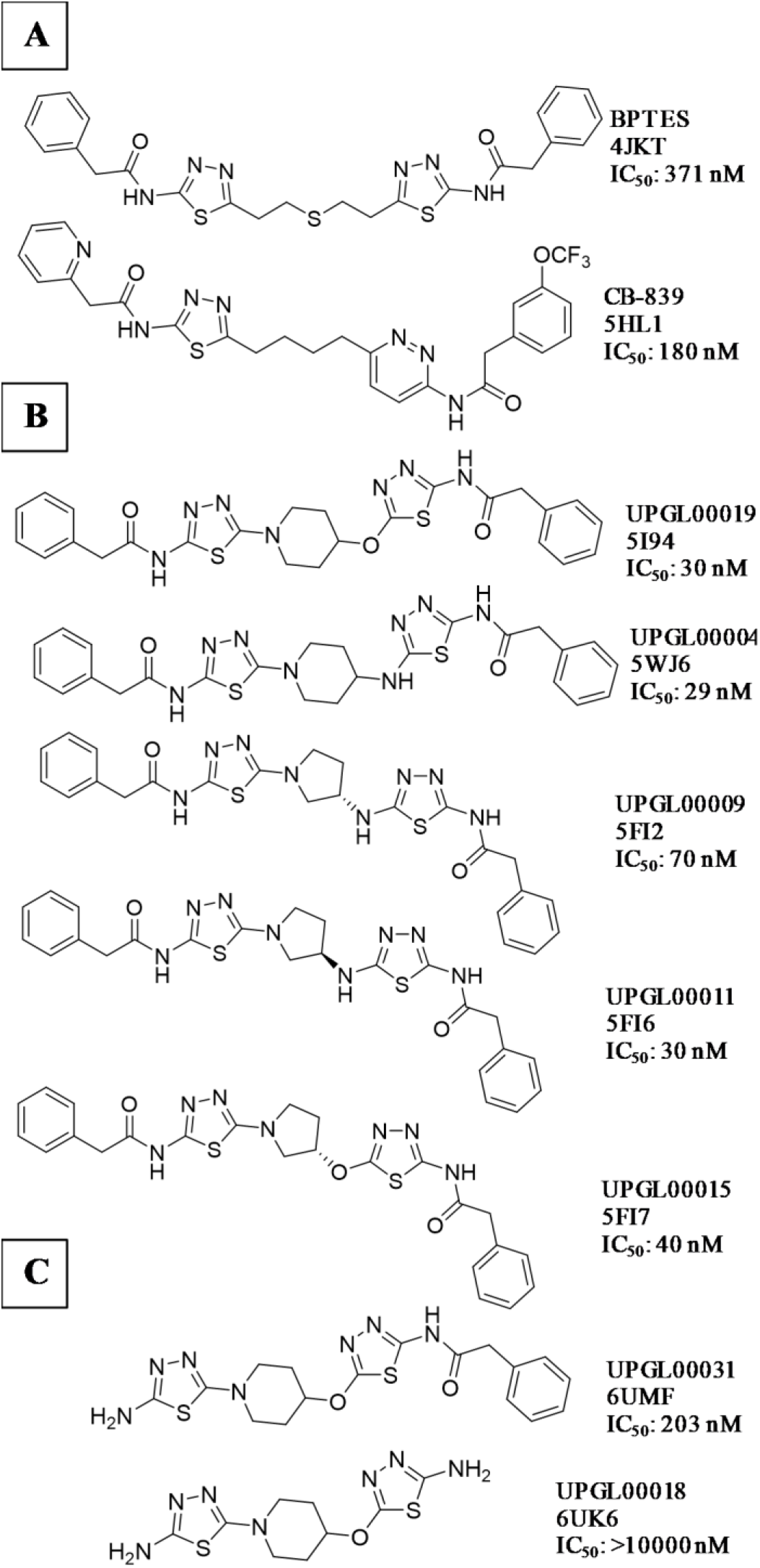
Allosteric inhibitors of GAC. A) BPTES, the parent member of the class of inhibitors, and CB-839, which is currently in clinical trials for various cancer indications. B) Five compounds from the UPGL series for which we had previously reported crystal structures. C) Two examples of the eleven UPGL series compounds co-crystallized with GAC in this study. PDB ID codes are shown for each compound. IC_50_ values reported are taken from (L. A. McDermott et al., 2016).

In the present study we aimed to further probe the molecular determinants responsible for the potency of the BPTES/CB-839 class of inhibitors and their mechanisms of action. First, we solved the X-ray crystal structures of eleven additional molecules from the UPGL series bound to GAC (Figs. 1C and S1). These structures highlight a set of highly conserved contacts between the central cores of the UPGL compounds and the protein, which are maintained regardless of drug potency. By making use of serial room temperature crystallography (Biel, Thompson, Cunningham, Corn, & Fraser, 2017; Fraser et al., 2011; Illava et al., 2021), we then obtained our first insight into what dictates potency differences for the BPTES/CB-839 class of inhibitors. Next, we used computational approaches together with recently developed inhibitor binding assays to complement our crystallographic analyses to help further define the chemical differences between weakly and strongly potent inhibitory molecules. These studies shed new light on the molecular basis for the range of inhibitory potencies exhibited by the BPTES/CB-839 class of compounds and the mechanism by which they inhibit enzymatic activity, thus helping to inform future efforts toward designing GAC inhibitors that combine improved potency with favorable pharmacological characteristics.

## Results

### X-ray crystal structures for GAC bound to the BPTES/CB-839 class of inhibitors show a conserved binding interaction despite differences in inhibitory potency

To obtain a better understanding of how the BPTES/CB-839 class of inhibitors bind to and inhibit GAC enzymatic activity, we solved the X-ray crystal structures of GAC complexed to eleven different inhibitors from the UPGL series of compounds (Figs. 1C and S1, Table S1). The crystals were cryo-cooled at high pressure (350 MPa) before placing them in the X-ray beam (Qingqiu Huang et al., 2018) to improve diffraction data. The structural analyses showed that each compound assumes a cup-like orientation within the helical interfaces between two GAC dimers. An example for GAC bound to compound UPGL00031 is shown in Fig. 2. Members of the UPGL series engage in a conserved hydrogen bonding network via their thiadiazole rings (or pyridazine rings in the case of UPGL00045) to the backbone atoms of Lys 320, Phe 322 and Leu 323 of GAC, and/or the hydroxyl hydrogen of Tyr 394, similar to what has been observed for BPTES and CB-839 (Qingqiu Huang et al., 2018). All of the compounds largely occupy the same region of space, with the sole outlier being UPGL00031, which shifts slightly in the binding site to enable its primary amine to form a hydrogen bond to the backbone carbonyl of Asn 324 while still maintaining the hydrogen bonding network shared across the UPGL series. The central cores of the UPGL-series of molecules take on a variety of conformations in order to project the thiadiazole rings into this hydrogen bonding network, leading to the hypothesis that correctly positioning the thiadiazole rings is a major requirement for binding.

**Figure 2:**
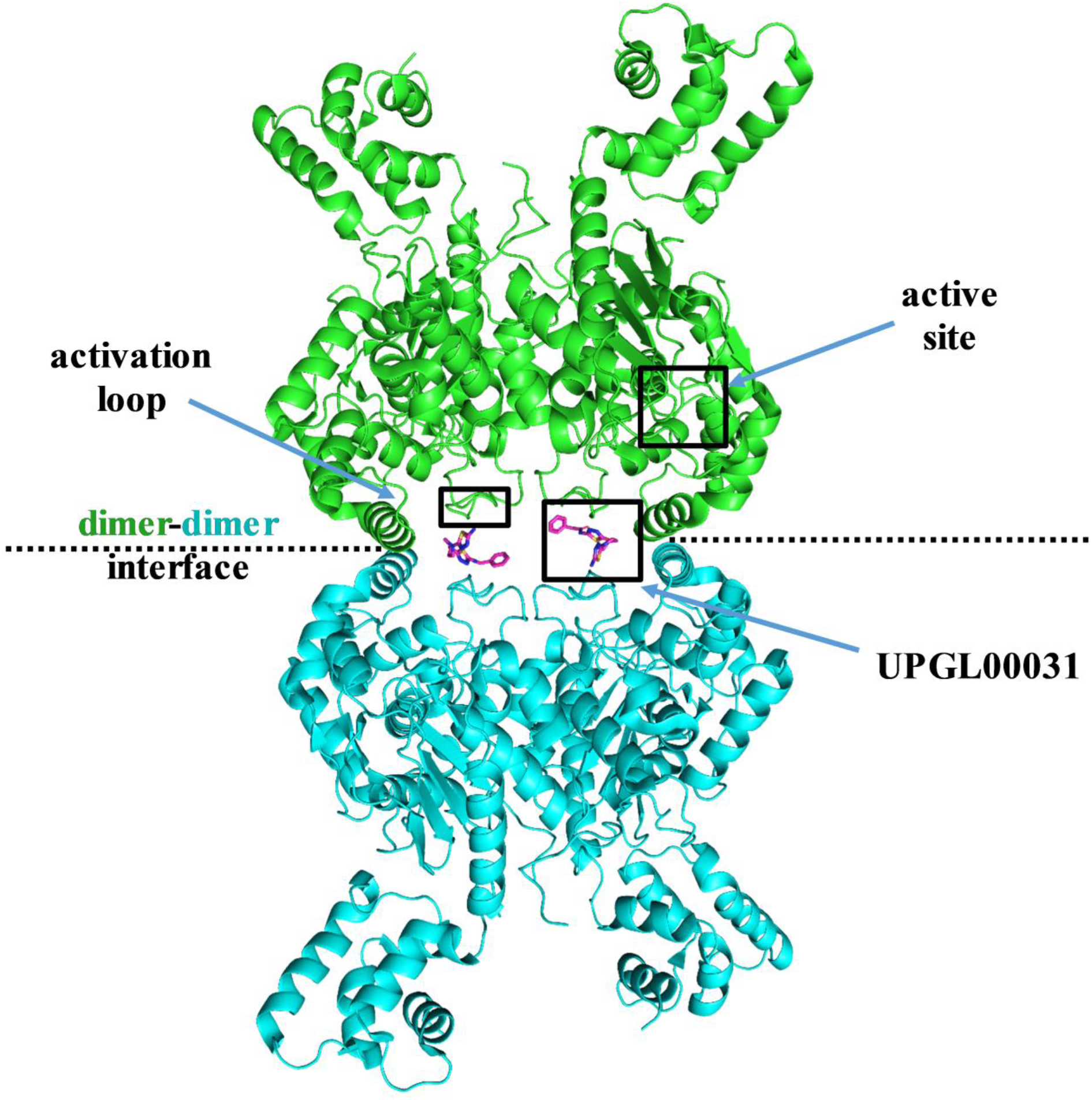
Cryo-cooled X-ray crystal structure of GAC bound to UPGL00031 (PDB ID 6UMF). GAC crystallized as a tetramer, with two dimers coming together to form a tetramer along the dotted line. Each GAC monomer has one catalytic site and one activation loop. The relative positions of the catalytic site, the activation loop, and the binding sites for the BPTES/CB-839 class of inhibitors are indicated. Bound UPGL00031 is shown in magenta.

As shown in Figs. 3A and 3B, even compounds of the UPGL series with vastly different potencies, e.g. UPGL00019 and UPGL00018, assume nearly identical orientations with the same hydrogen bonding network to GAC. However, the electron density for the terminal rings of the compounds and for several GAC residues in the activation loop could not be fully resolved in the X-ray crystal structures for the different GAC-inhibitor complexes (such as UPGL00019; Fig. 3A). The B-factors, which describe the degree of atomic motion in a crystal structure, for the terminal rings of UPGL00019 and neighboring GAC residues are relatively high (colored red and orange), despite the low B-factors (blue and green colors) for the core of the molecule and most of the residues in the activation loop. This trend is consistent across all of the co-crystal structures of GAC and the UPGL compounds.

**Figure 3:**
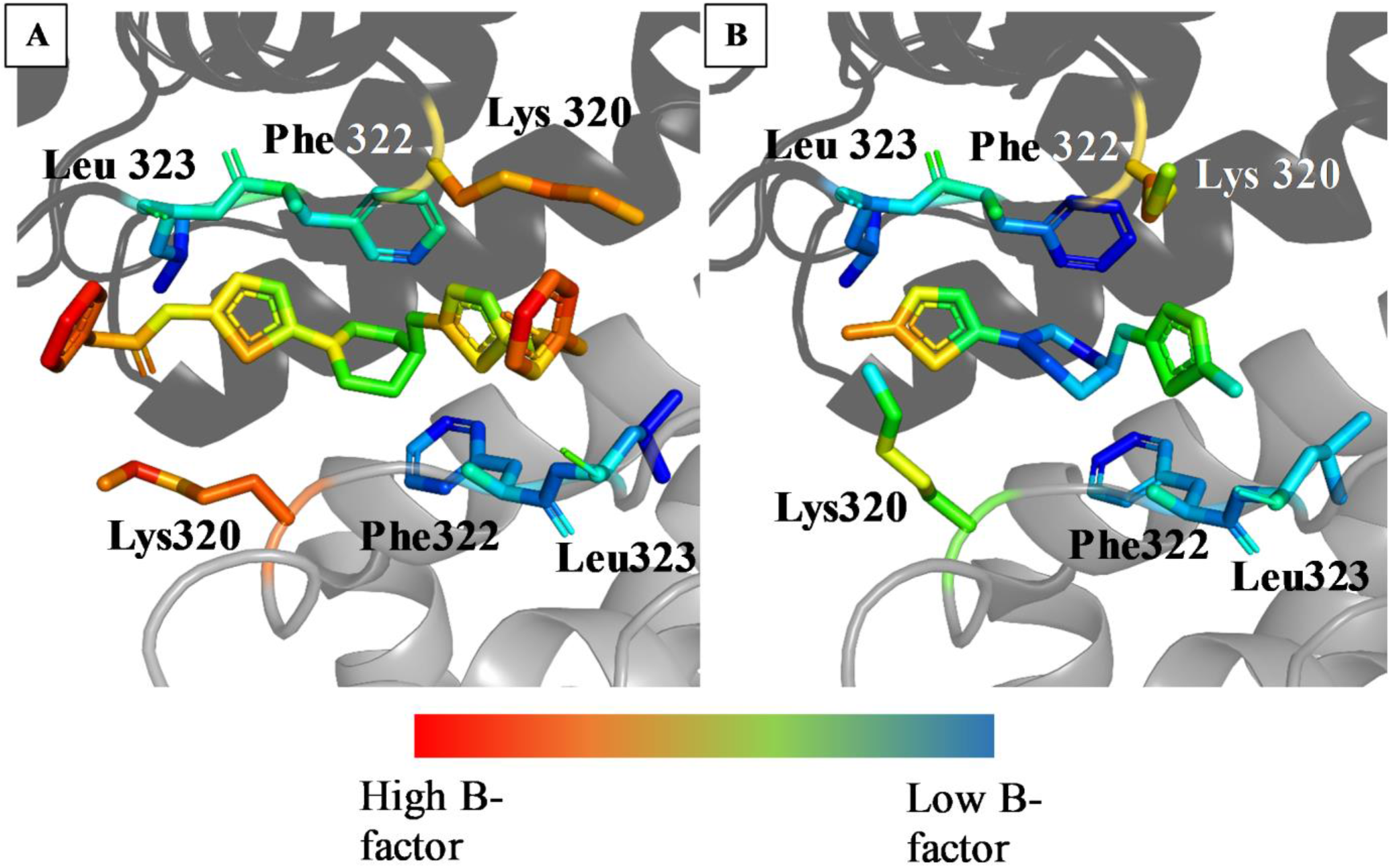
Cryo-cooled X-ray crystal structures show that BPTES/CB-839-class molecules bind to GAC in a similar fashion regardless of inhibitory potency. A) Cryo-cooled X-ray crystal structure of UPGL00019 bound to the activation loop of GAC. Each UPGL00019 molecule is bound to two adjacent monomers, which are presented as ribbon and colored as different shades of gray. The drug molecule and residues in the activation loop that engage in hydrogen bonds with the drug are colored by B-factors. Blue and green B-factor coloration suggest regions of little movement, while red and orange colors suggest regions of greater movement in the crystal structure. Most of the activation loop, and the core of the inhibitor, have low B-factors, but the GAC residues proximal to Lys 320, and the terminal phenyl rings of the inhibitor, are all more highly mobile. The other crystal structures in the series show similar trends. B) Cryo-cooled X-ray crystal structure of UPGL00018 bound to GAC (PDB ID 6UK6) colored as in (A). UPGL00018 occupies a nearly identical region of space in GAC to UPGL00019, despite enormous potency differences.

### Serial room temperature X-ray crystallography provides an insight into the basis for the marked differences in potency exhibited by two members of the BPTES/CB-839 class of inhibitors

Given that the traditional cryo-cooled crystallography described above could not differentiate between drugs with different potencies, we turned to serial X-ray crystallography, which collects data from dozens of individual crystals at room temperature to achieve a high-resolution structure, and offers the potential to reveal dynamic ligand-binding states not detected when using cryogenic methods (Fraser et al., 2011). Crystallization and data collection of complexes between GAC and either UPGL00004 (a potent inhibitor) or BPTES (a less potent inhibitor) were performed at room temperature and the diffraction data for at least sixty crystals of each complex were analyzed to determine their structures (see Table S2 for crystallization parameters).

The serial room temperature crystal structure that was solved for the GAC-UPGL00004 complex was largely similar to the corresponding high-pressure cryo-cooled structure, with the two inhibitor molecules bound to the GAC tetramer exhibiting the same cup-shaped orientation routinely observed in the cryo-cooled structure for this complex. Figs. 4A and 4B show examples for one of the two bound UPGL0004 molecules as observed in the room temperature (4A) and cryo-cooled (4B) crystal structures. This indicates that the high-pressure cryo-cooling was not distorting the protein significantly. However, the serial room temperature structure for the GAC-BPTES complex shows that one of the two BPTES molecules assumes a more extended orientation (Fig. 4C) compared to UPGL00004, whereas in the cryo-cooled co-crystal structure of GAC bound to BPTES, each of the two bound BPTES molecules adopt an orientation similar to UPGL00004 (Fig. 4D). Moreover, in the room temperature structure for the GAC-BPTES complex, the thiadiazole ring within the more linear end of the extended BPTES molecule is shifted away from its hydrogen bonding partners, the backbone atoms of Phe 322 and Leu 323. This disruption of hydrogen bonds between GAC and the thiadiazole ring in BPTES, compared to the same ring in UPGL00004, might be the basis for BPTES having a weaker binding affinity and lower potency with GAC.

**Figure 4:**
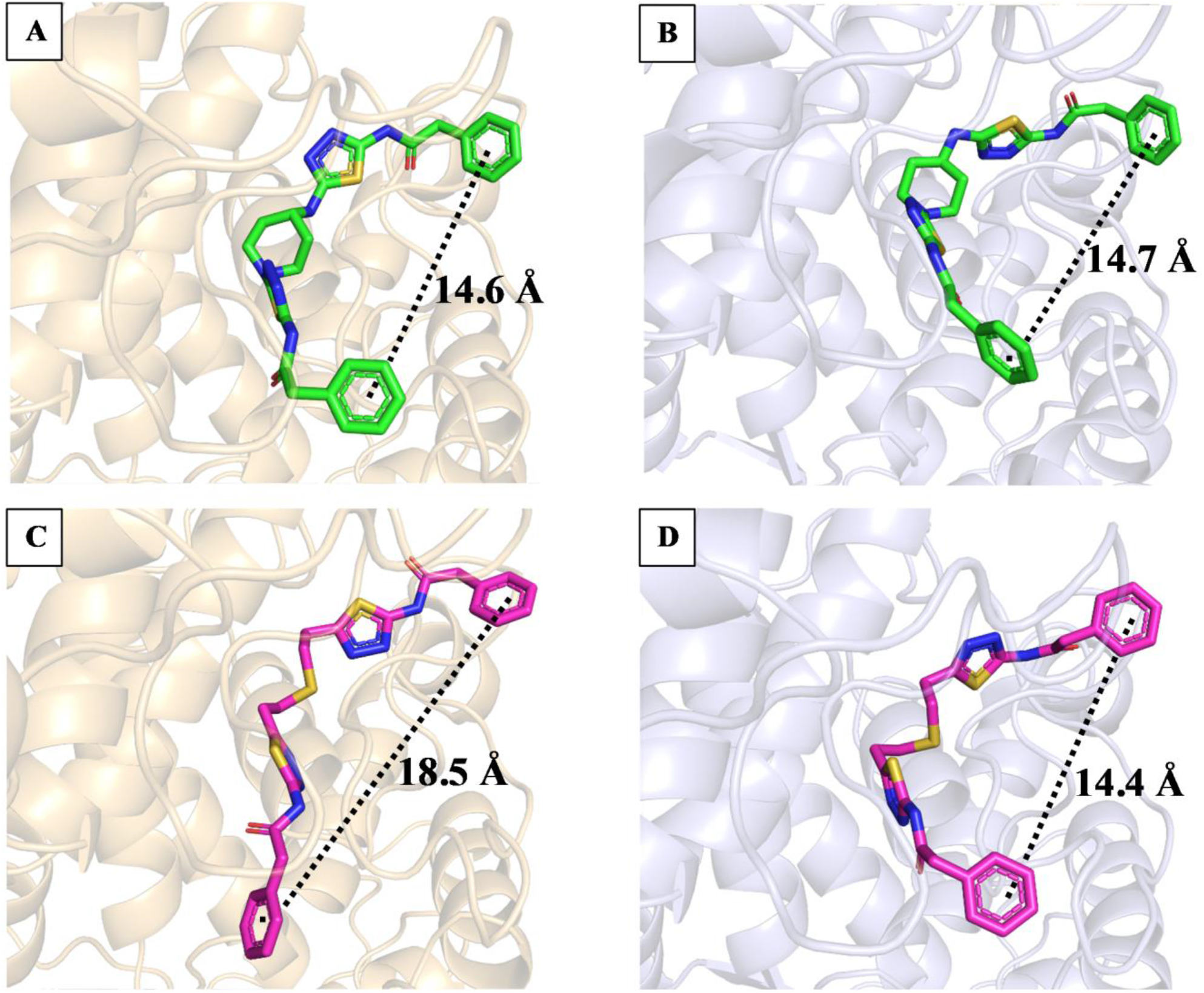
Comparison between the serial room temperature and cryo-cooled X-ray crystal structures of inhibitor-bound GAC complexes. A) Serial room temperature crystal structure showing one of the two UPGL00004 molecules (green) bound to GAC (light orange). The two molecules of UPGL0004 bound to GAC adopt the expected cup-shaped structure. B) Cryo-cooled crystal structure showing one of the two UPGL00004 molecules (green, PDB ID 5WJ6) bound to GAC (light blue). The structure of GAC, and conformation of UPGL00004, are nearly identical to the serial structure in (A). C) Serial room temperature crystal structure showing that one of the two BPTES molecules (magenta) bound to GAC (light orange) adopts an extended, semi-linear conformation. D) Cryo-cooled crystal structure of one of the two BPTES molecules (magenta, PDB ID 4JKT) bound to GAC (light blue). Unlike in the structure in (C), both molecules of BPTES bound to GAC adopt the expected cup-like conformation.

### QSAR analysis highlights the importance of the terminal rings in determining the potency of the BPTES/CB-839 class of inhibitors

To supplement our understanding of inhibitory potency, we developed a QSAR (quantitative structure/activity relationship) model. This model is based exclusively on the chemical structures of the BPTES/CB-839 class of inhibitors, and therefore offers an independent approach free of any structural bias that might arise from the co-crystallization process with GAC. We calculated electron density-derived properties of ∼1000 BPTES/CB-839-class inhibitors (C.M. Breneman et al., 2003; Sukumar & Breneman, 2007; Whitehead, Breneman, Sukumar, & Ryan, 2003) and used unbiased feature selection and kernel partial least squares (KPLS) regression to predict their inhibitory potency. Fig. 5A shows that for the test data set, predicted inhibitory potency largely mirrored the experimentally determined inhibitory potency for the various compounds (training set data is presented in Fig. S2A). Twenty-nine properties highly correlated with inhibitory potency were retained in the trained model following unbiased feature selection (Table S3). Y-scrambled models in which inhibitor potency values were randomly reassigned to each compound were entirely non-predictive (Figs. S2B-S2D), supporting the accuracy of the models for predicting inhibitory potency, and demonstrating that the KPLS model was learning valuable chemical information.

**Figure 5:**
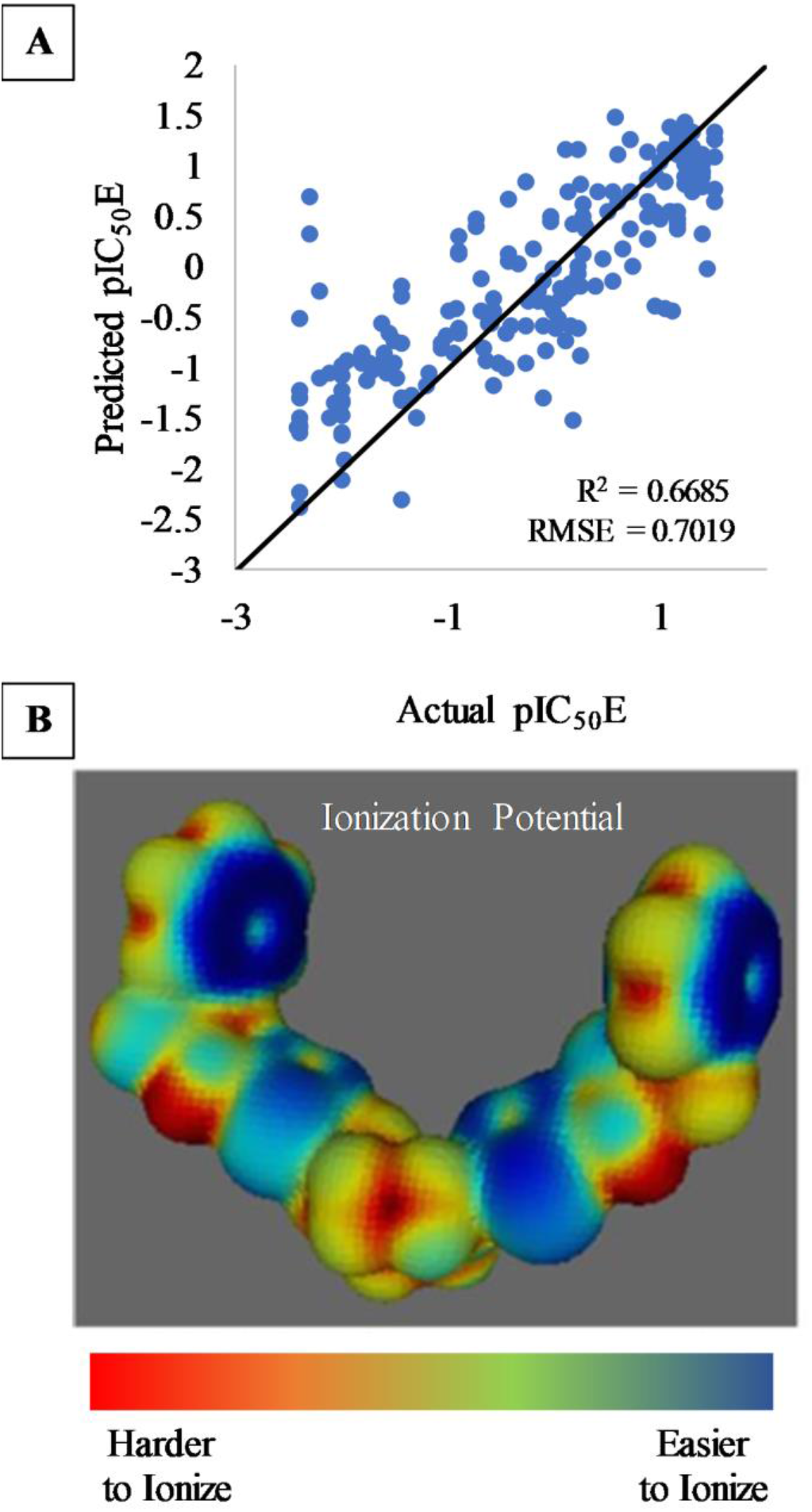
QSAR Modeling of the BPTES/CB-839 class of inhibitors. A) Test-set results for the QSAR model. The model has generally good accuracy, although it is prone to predicting high pIC_50_E values for some of the least inhibitory molecules (i.e. false positives). B) PIP (Politzer’s average local ionization potential) colored surface of UPGL00019. Blue regions are more easily ionizable, while red regions resist ionization. Coloration is scaled between the minimum and maximum PIP value for the molecule. The terminal rings, and thiadiazole rings, are the most easily ionized region of the molecule.

Among the twenty-nine descriptors used in the KPLS model were six descriptors describing relatively low values of the electrostatic potential (EP) and Politzer’s average local ionization energy (PIP) (Politzer, Bulat, Burgess, Baldwin, & Murray, 2012). All but one of these descriptors (FPIP3) were positively correlated with inhibitory potency (Table S3, positively correlated descriptors are colored green). Thus, we examined the PIP or EP surface maps of the different molecules. For the case of UPGL00019 and other compounds with similar potency, the terminal groups and thiadiazole rings had the largest surface areas with low PIP (i.e. highly ionizable surface area, Figs. 5B, S3). Similar results were observed for EP maps. This supported the indications from the structural analyses of GAC-inhibitor complexes that the hydrogen bonding network between the thiadiazole rings and protein backbone was a major source of inhibitor binding affinity. It also suggested that the terminal groups of the BPTES/CB-839 class of compounds are in some way exerting a significant influence on their inhibitory capability despite not being captured in the X-ray structures.

### Fluorescence spectroscopic read-outs for inhibitor binding to GAC support the role of the terminal rings in determining inhibitor potency

Recently, we developed two fluorescent spectroscopic assays to examine the coupling between inhibitor binding and conformational changes occurring either at the activation loop or the substrate-binding site of GAC (Li et al., 2019; Stalnecker, Erickson, & Cerione, 2017). These read-outs made use of GAC mutants in which a tryptophan replaced either a phenylalanine within the activation loop close to where the BPTES/CB-839 class of inhibitors bind (GAC (F322W)), or a tyrosine at the substrate-binding site (GAC (Y466W)) (Li et al., 2019; Stalnecker et al., 2017). To test the suggestion from our modeling efforts that the terminal groups of the BPTES/CB-839 class of inhibitors contribute to their ability to bind and affect GAC catalytic activity, we examined a subset of the UPGL series with identical molecular structure at the centers, but which differ in the number (but not the structure) of their terminal groups. These compounds were UPGL00019 which has two terminal phenyl rings (IC_50_ = 30 nM), UPGL00031 with a single terminal phenyl ring (IC_50_ = 200 nM), and UPGL00018 that lacks terminal groups (IC_50_ > 10,000 nM). We first examined their ability to alter the conformation of the activation loop, i.e. the site where the compounds bind to GAC (Li et al., 2019; Stalnecker et al., 2017). The GAC (F322W) mutant was treated with each of the three inhibitors, or with DMSO as a negative control, and the fluorescent signal of Trp 322 was monitored (Figs. 6A, 6B, S4A, and S4B). The binding of both UPGL00019 and UPGL00031 resulted in a quenching of Trp 322 fluorescence, while UPGL00018 was largely ineffective at micromolar concentrations. We then treated the GAC (Y466W) mutant with each molecule to monitor conformational changes within the substrate-binding site (Li et al., 2019; Stalnecker et al., 2017). UPGL00019 and UPGL00031 were again effective at causing a quenching of Trp 466 fluorescence, whereas UPGL00018 was markedly less effective (Figs. 6C, 6D, S4C, and S4D). These results indicate that the presence of at least one terminal group on the molecule can either help inhibitor binding and/or alter the activation loop conformation in a manner that blocks its communication with the active site, which in turn inhibits catalytic activity.

**Figure 6:**
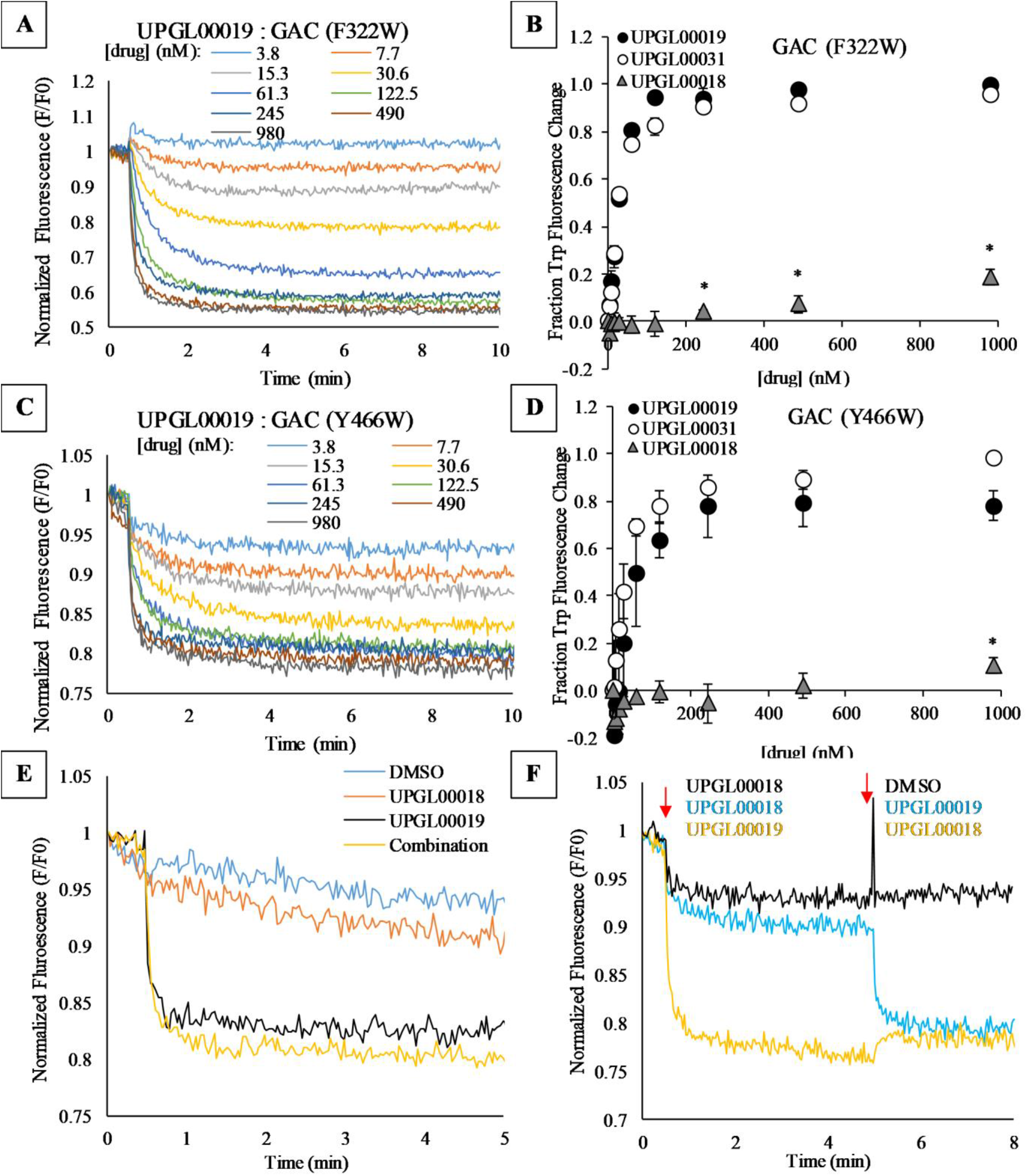
Analysis of the binding mechanism between GAC and the BPTES/CB-839 class of inhibitors. A) Real-time tryptophan fluorescence emission signal (λ_ex_ = 285nm, λ_em_ = 340nm) of 100 nM GAC (F322W) is quenched upon the addition of UPGL00019 at 30 seconds. The signal was normalized to the initial fluorescence (F_0_). B) The equilibrium fluorescence from panel A was plotted as a function of drug concentration for different UPGL-series inhibitors. While both UPGL00019 (black circles, K_d_ = 54.5 nM) and UPGL00031 (white circles, K_d_ = 34.4 nM) are able to strongly quench the tryptophan fluorescent signal, UPGL00018 binds weakly (gray triangles, K_d_ > 1,000 nM), but shows statistically significant quenching of GAC (Y322W) at concentrations as low as 240 nM. C) Time-dependent tryptophan fluorescence quenching of 100 nM GAC (Y466W) by UPGL00019. D) Normalized fluorescence quenching of GAC (Y466W) by UPGL00019 (black circles), UPGL00031 (white circles) and UPGL00018 (gray triangles). UPGL00019 (K_d_ = 28.7 nM) and UPGL00031 K_d_ = 27.2 nM) are each able to strongly quench the tryptophan fluorescent signal. UPGL00018 (gray triangles, K_d_ > 1,000 nM) has no statistically significant effect on tryptophan fluorescence until the highest tested concentration (980 nM), where the effect is minimal. Statistical significance is shown only for UPGL00018 data points. E) Binding assays for 100 nM GAC (Y466W) and 1 μM UPGL00018 and/or 1 μM UPGL00019, when the inhibitor molecules are added simultaneously to the enzyme. The quenching caused by an equimolar amount of both drugs together is identical to that caused by UPGL00019 alone, showing that UPGL00018 is unable to compete away UPGL00019. F) Binding assays for 100 nM GAC (Y466W) and 1 μM UPGL00018 and 1 μM UPGL00019 added sequentially. The drug added at each injection point for each curve is indicated on the plot. Order of addition does not affect the total level of Trp quenching, suggesting that kinetic variables (on-rate and off-rate) do not account for the inability of UPGL00018 to compete with UPGL00019 in these assays. Data shown in panels (A), (C), (E), and (F) are representative of three separate experiments.

We then examined if the inability of UPGL00018 to cause a detectable quenching of Trp 466 fluorescence was due to a significantly weaker binding affinity for GAC compared to UPGL00019, or if UPGL00018 is capable of binding to the enzyme with high affinity but is unable to induce the necessary conformational change to quench the tryptophan fluorescence emission. In one set of experiments, the GAC (Y466W) mutant was treated simultaneously with 1 μM UPGL00018 and 1 μM UPGL00019. As shown in Fig. 6E, the resultant quenching of Trp 466 fluorescence was nearly identical to that with UPGL00019 treatment alone. The same was true when UPGL00018 was added first to GAC followed by UPGL00019 (Fig. 6F). The inability of 1 μM UPGL00018 to block the binding of 1 μM UPGL00019 to GAC(Y466W) and its accompanying quenching of Trp 466 fluorescence indicates that it binds with a significantly weaker affinity compared to UPGL00019. Therefore, the terminal groups of the BPTES/CB-839 class of inhibitors appear to be important for their GAC-binding affinity and correspondingly, for their inhibitory capability.

### Lysine 320 is essential for the binding of the BPTES/CB-839 class of inhibitors to GAC

Because the terminal groups of the BPTES/CB-839-class inhibitors such as UPGL00019 are essential for high affinity binding to GAC, we were interested in determining how these rings interact with GAC. In some of the co-crystal structures for GAC bound to the BPTES/CB-839 class of molecules (e.g. 5HL1), Lys 320 projects toward at least one terminal ring of the bound inhibitor. Moreover, Lys 320 plays an essential role in catalysis, as substituting an alanine for the lysine residue at this position (GAC (K320A)) resulted in a constitutively active enzyme (Li et al., 2019). We prepared the double-mutant GAC (K320A, Y466W) to analyze the binding of two of the most potent compounds in the UPGL series, UPGL0004 and UPGL00019, by monitoring the changes in the fluorescence of Trp 466. The lysine to alanine mutation resulted in a striking reduction in the binding affinity (i.e. apparent K_d_ values) of these compounds to the enzyme (Table 1). We also examined inhibitor binding to the double-mutants GAC (R317A, Y466W) and GAC (F318A, Y466W), since Arg 317 and Phe 318 also project to be near the terminal rings of the inhibitor molecules. These alanine-substituted mutants however were fully capable of binding to each of the inhibitors (Table 1).

**Table 1:**
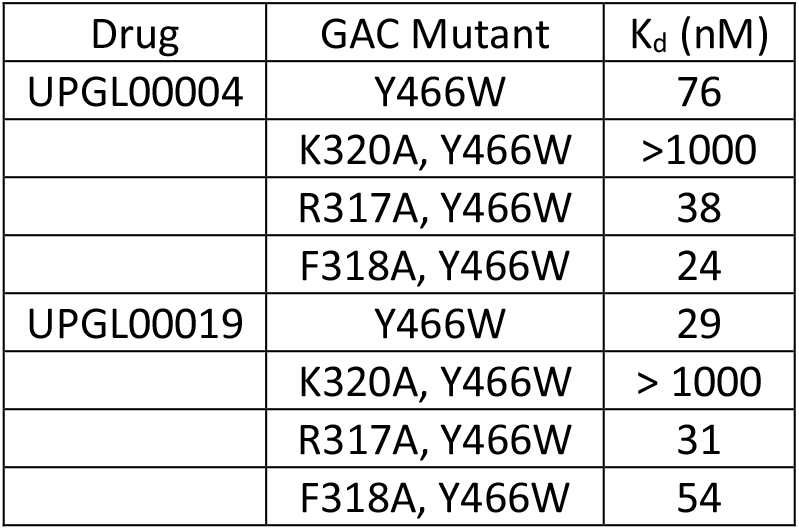
Apparent K_d_ values for UPGL00004 or UPGL00019 binding to different GAC mutants.

| Drug | GAC Mutant | $K_d$ (nM) |
| --- | --- | --- |
| UPGL00004 | Y466W | 76 |
|  | K320A, Y466W | >1000 |
|  | R317A, Y466W | 38 |
|  | F318A, Y466W | 24 |
| UPGL00019 | Y466W | 29 |
|  | K320A, Y466W | > 1000 |
|  | R317A, Y466W | 31 |
|  | F318A, Y466W | 54 |

## Discussion

GAC has garnered significant attention as a potential cancer target, with considerable effort spent on studying the BPTES/CB-839 class of compounds. However, thus far, no GAC inhibitor has been approved for cancer treatment. A major shortcoming for the further development of the BPTES/CB-839 family of molecules as drug candidates stems from questions regarding what dictates their potency. Our analyses of 11 new X-ray crystal structures for GAC complexed to members of the UPGL series of the BPTES/CB-839 compound family of varying inhibitory potency, using crystals obtained under cryo-cooled conditions, showed that the binding contacts for the different inhibitors were largely conserved. Thus, despite these extensive crystallization efforts, the molecular determinants that dictate potency for this class of GAC allosteric inhibitors were not evident.

To gain further insight, we took advantage of serial room temperature crystallography to examine two allosteric inhibitors of GAC with different potencies, BPTES and UPGL0004. The room temperature X-ray crystal structure for GAC complexed to BPTES showed two distinct poses, with one of the BPTES molecules exhibiting an extended conformation, and the other more closely resembling the prior cup-shaped structural images of this inhibitor. By contrast, *both* molecules of UPGL00004 bound to GAC in the room temperature X-ray structures adopted the more typical cup-shaped orientation within the binding site. This supports the idea that UPGL00004 is held rigidly in the allosteric binding site of GAC, while BPTES, even when bound to the enzyme, has a significant degree of conformational flexibility. Moreover, our results suggest that proper positioning of the thiadiazole rings of either compound within the hydrogen bonding network of the allosteric binding site is one of the key determinants of inhibitory potency, and that the inability to maintain the proper positioning of these rings is one reason BPTES is less potent than UPGL00004.

Our QSAR model had an R^2^ of 0.67 for the blind test set, and compares favorably with earlier work by Amin and coworkers, who reported a QSAR model for this class of GAC inhibitors with a narrower dataset of only 40 compounds (Amin, Adhikari, Gayen, & Jha, 2017). Our model made use of six descriptors indicating the importance of highly polarizable, electronegative surfaces, and five of these descriptors (17% of all descriptors in the model) were positively correlated with inhibitory potency. Thiadiazole rings, which both cryo-cooled and room temperature crystallography suggested were a key to binding affinity, have highly polarizable, electronegative surface regions. However, the terminal rings of the most potent UPGL series molecules also have these characteristics, suggesting that they are also important for binding to GAC. This supports our earlier work suggesting that inhibitory potency might be influenced by the combined Van der Waals volume of the terminal groups of these inhibitors (L. McDermott et al., 2019) and is consistent with SAR data showing that for the BPTES/CB-839 class of molecules, the most potent compounds tend to either have aromatic rings (e.g. benzene or pyridine rings) or electron-donating pseudo-aromatic groups such as cyclopropane as terminal groups (M. I. Gross et al., 2014; William P Katt et al., 2017; Shukla et al., 2012; Xu et al., 2019). Moreover, fluorescent binding assays showed that at least one terminal ring is necessary for an inhibitor molecule to bind GAC with high affinity. This is evident as both UPGL00019 and UPGL00031 exhibit nearly identical binding profiles despite having different number of rings. However, the symmetric presence of the second terminal ring in UPGL00019 appears to help transmit the binding affinity from the activation loop to effective inhibition of catalytic activity in the active site, resulting in a ∼10-fold increase in the IC_50_ value compared to UPGL00031.

### Implications for the rational design of new inhibitor molecules

A key structural feature of the BPTES/CB-839 class of compounds necessary for tight binding and potent inhibition of GAC activity is the thiadiazole-centered hydrogen bonding network between the core of these inhibitor compounds and the enzyme, which is best maintained by a cup-shaped molecule. Indeed, a recently reported series of BPTES-derived molecules used macrocyclization to successfully stabilize this orientation and achieve low nanomolar potency, albeit with generally poor pharmacological properties (Xu et al., 2021). We also found that Lys 320 likely interacts with the terminal groups of BPTES/CB-839 class inhibitors and is essential for their ability to bind to GAC. This is further supported by the findings of Ferreira et al. that showed mouse GAC (K325A) (equivalent to human GAC (K320A)) is resistant to inhibition by BPTES at concentrations as high as 10 μM (Ferreira et al., 2013), which is similar to the concentration required for UPGL00018, a compound that lacks terminal rings, to elicit a minimal inhibitory effect (∼20% inhibition at 10 μM).

Overall, our findings support a mechanism in which the terminal rings of most BPTES/CB-839 class inhibitors initially undergo a dynamic and/or transient association with Lys 320 within the activation loop of GAC, which precedes a more stable, high affinity interaction involving a hydrogen bonding network between the thiadiazole rings of these molecules and the enzyme. Our data shows that a single terminal ring may be sufficient to engage this mechanism, as UPGL00031 and UPGL00019 have largely similar abilities to bind to GAC. The relatively high affinity of the enzyme for UPGL00031 might be because the terminal amine of this compound comes within 4 Å of the sidechain of Lys 320, potentially enabling it to engage in a hydrogen bond that compensates for the absence of a terminal ring. Although UPGL00031 is less potent than UPGL00019 in catalytic assays, and of similar potency to CB-839 (Fig. 1), it has a much higher LipE (4.42 for UPGL00031 vs. 3.36 for UPGL00019 or 1.99 for CB-839) (L. A. McDermott et al., 2016). LipE, which combines inhibitory potency and logP into a single number, is increasingly being recognized as a more accurate predictive factor for eventual clinical success than inhibitory potency alone (Freeman-Cook et al., 2012; Jabeen, Pleban, Rinner, Chiba, & Ecker, 2012). Thus, while new compounds that either enforce the cup-shaped orientation, or have groups specifically designed to interact with Lys 320, might be capable of higher potency compared to current molecules, there may also be a significant benefit in further optimizing compounds with a single terminal group, which have thus far been comparatively poorly studied.

## Conclusions

A major objective of these studies was to better define the molecular basis for the binding affinities and inhibitory potencies of the BPTES/CB-839 class of allosteric inhibitors for GAC, the key metabolic enzyme responsible for satisfying the glutamine addiction of a number of cancers. Using serial room temperature crystallography, we now describe how two members of the family with markedly different potencies exhibit distinct orientations as they interact with GAC. We further show that their inhibitory mechanism requires interaction with Lys 320 within the activation loop of GAC. These structural and mechanistic details will hopefully aid in the future development of the next generation of BPTES/CB-839-class inhibitors and help bring these molecules into the clinic.

## Materials and Methods

All small molecules were prepared as previously described (L. A. McDermott et al., 2016). Common chemicals and other consumables were obtained from Thermo Fisher Scientific (Waltham, MA). The IC_50_ values reported in Figs. 1 and S3 were taken from (L. A. McDermott et al., 2016). In that study, the compounds were assayed against the same preparation of recombinant GAC (50 nM), and total glutamine hydrolysis was determined via a coupled glutamate dehydrogenase assay.

### High pressure cryo-cooled crystal structures

Protein purification and crystallization were carried out as described previously (Qingqiu Huang et al., 2018; L. A. McDermott et al., 2016). (Qingqiu Huang et al., 2018; L. A. McDermott et al., 2016). Briefly, the indicated inhibitor was mixed with human GAC protein at a mole ratio of 4:1 and incubated on ice for one hour. Crystals were grown at 20°C in 10% PEG6000 (w/v), 1 M LiCl, and 0.1 M Tris-HCl, pH 8.5. Generally, crystals were observed within 24 hours, and reached a size of 100 × 100 × 200 μm^3^ after 7 days. The crystals were high-pressure cryo-cooled at 350 MPa for 30 min to reduce lattice disorder prior to data collection (Englich et al., 2011; Q. Huang et al., 2016). Diffraction data were collected at 100K at the CHESS A1 station. The diffraction data were processed using the HKL package (Otwinowski & Minor, 1997). Statistics of data collection and processing are summarized in Table S1.

The crystal structures were solved by molecular replacement using human apo GAC (PDB ID 5D3O) as a search model (Li et al., 2016). Model building was performed using COOT (Emsley & Cowtan, 2004), and refinement was performed using Phenix refine (Adams et al., 2010). Statistics of structure refinement are summarized in Table S1.

Both human and mouse GAC, which we have found to be catalytically identical, have been used in these studies. For simplicity, all residue numbering throughout the manuscript is based on the human GAC sequence, except when describing the methods for preparing mutants of mouse GAC below.

### Serial room temperature crystallography

Protein purification and crystallization were carried out using methods previously described (Qingqiu Huang et al., 2018; L. A. McDermott et al., 2016). Briefly, solutions of 20 mg/mL GAC (in 150 mM NaCl and 50 mM Tris-HCl, pH 7.5) and 30 mM inhibitor (BPTES or UPGL00004 in DMSO) were prepared. The protein-inhibitor complexes were formed by mixing 95 μL of the GAC solution and 5 μL of the inhibitor solution, yielding a mole ratio of 1:4, and then incubating the mixture on ice for 1 h. Crystals were grown at 20 °C by the hanging drop vapor diffusion method in crystallization trays. Typically, 1 μl of the complex solution was mixed with 1 μl of a reservoir solution consisting of 10% PEG6000 (w/v), 1.0 M LiCl, and 0.1 M Tris-HCl buffer (pH 8.5). Crystals were observed within 24 h, reaching an average size of 100 × 100 × 200 μm^3^ after 7 days. The crystals were transferred onto chips (sample support) mounted in crystal caps provided by MiTeGen (Ithaca, NY). Approximately 15-20 crystals were harvested per chip. The crystals on the chip were moved into a humidified glovebox (humidity >97%) (MiTeGen) and a vacuum was applied to remove excess liquid (from crystal harvesting) before sealing the chip with a thin transparent film (MiTeGen). In the beam (ID7B2 station at CHESS), the chips were raster scanned in 20 μm steps and 5° of oscillation data was collected. Each step of the raster scanning was completed in 0.75 s – 0.5 s and 0.25 ms for data acquisition (25 frames, 0.2° and 10 ms/frame) – corresponding to a 1.3 Hz raster rate. Individual oscillation frame sets were processed with XDS, and scaled and merged together with XSCALE (Kabsch, 2010b, 2010a). The detailed processing and filtering routine using XSCALE_ISOCLUSTER (Brehm & Diederichs, 2014) has been previously described (Wierman et al., 2019). Phasing and molecular replacement (using PDB ID: 5WJ6 as the phasing model) were performed using PHASER and phenix.refine in PHENIX, respectively (Adams et al., 2010; Liebschner et al., 2019).

### Comparison of crystal structures

All visualization was performed in PyMol (Schrodinger LLC, 2010). Crystal structures were aligned using the “cealign” command targeting the whole protein structure.

### Computational chemistry

Molecules for QSAR modeling were taken from several sources (Bennett et al., 2014; Bhavar, Vakkalanka, Viswanadha, Swaroop, & Babu, 2015a, 2015b; Cianchetta, Lemieux, Cao, Ding, & Ye, 2015; Di Francesco et al., 2016; L. A. McDermott et al., 2016; Shukla et al., 2012; Zimmermann et al., 2016), and data were entered using the ChemFinder plugin for Excel. Molecular structures were converted to SMILES strings, and the SMILES strings were used as input in the RECON software package for descriptor generation (Curt M Breneman et al., 2003; Sukumar & Breneman, 2007). Because multiple datasets were combined for modeling purposes, IC_50_E (IC_50_ effective) values were defined as the source’s reported IC_50_ for the molecule divided by the IC_50_ for BPTES according to that source. Where compound potency was reported as classification data (Bhavar et al., 2015a, 2015b; Cianchetta et al., 2015; Di Francesco et al., 2016), compound IC_50_ was entered as the highest reported IC_50_ for the classification range. These IC_50_E values were then converted to pIC_50_E values for use in building the model.

A random test and training set were generated for the molecules by assigning each molecule a random value (0 to 1) in Excel. Molecules with a value ≤ 0.2 were assigned to the test set (203 molecules), all other molecules were assigned to the training set (722 molecules). The models were built using tools from the Rensselaer Exploratory Center for Cheminformatics Research (RECCR): the RECCR Online Modeling System (ROMS) (Zhen, Potta, A. Lanzillo, Rege, & M. Breneman, 2017), and the SVR-Based Online Learning Equipment (SOLE) (Zhen et al., 2017). A kernel partial least squares (KPLS) model was trained in ROMS using feature selection. This included removal of descriptors which were 4-sigma outliers (descriptors which had individual values lying more than 4 standard deviations from the mean of all values of that descriptor), and removal of descriptors that were ≥ 90% correlated to other descriptors included in the model. KPLS models were prepared using bootstrapping (100 rounds of bootstrapping with 100 molecules withheld as an internal test set). The models used 5 latent variables, and the kernel used a sigma value of 10. LS-SVR models were trained in SOLE using feature selection (85% mutual correlation threshold), 10-fold cross validation with 10% of the dataset reserved for this purpose, a linear kernel, and default gamma and sigma parameters, and showed largely similar results to the KPLS models.

To conduct y-scrambling, the assorted pIC_50_E values for the molecules were reassigned randomly to each molecule. The data, with scrambled pIC_50_E values, was then modeled in exactly the same way as the unscrambled data. This procedure was repeated 10 times, each with a different random assignment of pIC_50_E values to the molecular input data.

Isosurfaces colored by either electrostatic potential (EP), or average local ionization energy (PIP) (Politzer et al., 2012), were prepared by calculation of the properties in Gaussian ‘09 (Frisch et al., 2013). The properties were then mapped to *ρ* (**r**) = 0.002 e bohr^-3^ isosurfaces using in-house codes developed by N. Sukumar, C. Breneman, and the RECCR team at Rensselaer Polytechnic Institute.

### Fluorescent tryptophan quenching assays

Mouse GAC (F327W) and GAC (Y471W), which correspond to human GAC (F322W) and GAC (Y466W), were prepared as previously described (Qingqiu Huang et al., 2018; Stalnecker et al., 2017), except that the protein was eluted in a higher salt buffer during FPLC purification (500 mM NaCl, 20 mM Tris-HCl, pH 8.5) to enhance protein stability. Assays were then conducted as previously described (Qingqiu Huang et al., 2018; Stalnecker et al., 2017). Briefly: 100 nM GAC (F327W) or GAC (Y471W) was solvated in 1 mL of 50 mM Tris-acetate, pH 8.5, with 0.1 mM EDTA and the indicated amount of inhibitor. Samples were stirred constantly while being held at 25 °C and were measured using a Varian Cary Eclipse fluorimeter in counting mode, with an excitation wavelength of 285 nm (5 nm bandpass) and an emission wavelength of 340 nm (20 nm bandpass). P-values were calculated using Student’s 2-tailed t-test from normalized fluorescence values.

## Acknowledgements

W.P.K, L.A.M., and R.A.C. conceived the study. S.K.M., Q.F., A.F., I.K., D.S., M.S., and E.A. conducted the serial crystallography experiments. Q.H. conducted the cryo-cooled crystallography experiments. T.T.N. and S.R. conducted the fluorescence quenching experiments. N.S., and W.P.K. conducted the QSAR experiments. L.A.M. provided the small molecules for study. W.P.K. and R.A.C. wrote the paper. All authors read and agreed upon the final manuscript before submission. We would also like to thank the staff at CHESS for their assistance with the crystallographic experiments, and the late Dr. Jon Erickson for many valuable conversations. This work was supported by grants to R.A.C. (GM122575, CA201402), L.A.M. (University of Pittsburgh Medical Center Competitive Medical Research Fund (CMRF)), CHESS (P30GM126166), and to N. Sukumar (Science & Engineering Research Board (SERB), India, grant# EMR/2016-002141). We also thank MiTeGen for supplying key resources for the serial X-ray crystallography experiments.

