## Supplemental Tables and Figures for "New Insights into the Mechanisms Used by Inhibitors Targeting Glutamine Metabolism in Cancer Cells"

### Supplemental Information:

| Inhibitor | UPGL-12 | UPGL-13 | UPGL-18 | UPGL-20 | UPGL-23 | UPGL-27 | UPGL-30 | UPGL-31 | UPGL-45 | UPGL-60 | UPGL-61 |
|---|---|---|---|---|---|---|---|---|---|---|---|
| **Data collection** | | | | | | | | | | | |
| Space group | P21 | P21 | P21 | P212121 | P212121 | P212121 | P21 | P21 | P21 | P212121 | P212121 |
| Cell dimensions | | | | | | | | | | | |
| a (Å) | 98.83 | 49.30 | 54.44 | 99.23 | 97.88 | 99.98 | 98.72 | 49.73 | 49.55 | 100.48 | 100.87 |
| b (Å) | 138.84 | 139.01 | 138.79 | 139.39 | 138.24 | 138.82 | 138.89 | 138.98 | 138.89 | 139.16 | 139.09 |
| c (Å) | 177.21 | 177.39 | 179.05 | 178.13 | 176.49 | 176.64 | 177.15 | 177.03 | 176.99 | 178.19 | 178.24 |
| β (°) | 90.03 | 93.41 | 89.99 | 90 | 90 | 90 | 89.98 | 93.91 | 93.91 | 90 | 90 |
| Resolution (Å) | 50-2.9 | 50-2.5 | 50-2.7 | 50-3.0 | 50-2.5 | 50-2.6 | 50-2.7 | 50-2.7 | 50-2.7 | 50-2.5 | 50-2.8 |
| | (2.95-2.90) | (2.54-2.50) | (2.75-2.70) | (3.05-3.0) | (2.54-2.50) | (2.64-2.60) | (2.75-2.70) | (2.75-2.7) | (2.75-2.70) | (2.54-2.50) | (2.85-2.80) |
| Unique reflections | 105300 | 78825 | 71136 | 50186 | 82779 | 71710 | 124737 | 63034 | 56593 | 85958 | 121885 |
| | (5072) | (3492) | (3135) | (2448) | (4060) | (3067) | (6102) | (2581) | (2566) | (3800) | (5554) |
| Redundancy | 3.4 (3.2) | 6.2 (5.8) | 3.4 (2.6) | 7.9 (7.4) | 6.6 (5.9) | 7.2 (6.3) | 3.3 (3.1) | 5.2 (4.4) | 3.2 (2.6) | 6.8 (6.0) | 3.7 (3.1) |
| Completeness (%) | 98.5(95.8) | 95.4(85.1) | 96.4(84.4) | 99.7(99.4) | 99.6(99.1) | 97.9(84.1) | 97.2(95.7) | 93.6 (77.3) | 85.8(76.4) | 98.2(88.6) | 99.2(91.2) |
| Average I/σI | 6.39(0.98) | 8.78(1.23) | 11.21(0.15) | 13.54(2.30) | 16.23(4.71) | 13.94(1.44) | 9.14(0.86) | 4.1 (0.45) | 15.64(2.80) | 15.72(1.59) | 8.89(1.05) |
| $R_{sym}$ | 0.257(0.0) | 0.119(0.512) | 0.138(0.0) | 0.227(0.0) | 0.199(0.491) | 0.20(0.0) | 0.169(0.0) | 0.237(0.684) | 0.110(0.484) | 0.134(0.0) | 0.156(0.888) |
| $R_{meas}$ | 0.293(0.0) | 0.129(0.558) | 0.163(0.0) | 0.234(0.0) | N/D | N/D | 0.185(0.0) | 0.282(0.934) | 0.131(0.597) | 0.144(0.0) | 0.173(0.0) |
| CC1/2 | 0.573 | 0.883 | 0.523 | 0.860 | N/D | N/D | 0.528 | 0.645 | 0.739 | 0.903 | 0.811 |
| CC* | 0.854 | 0.968 | 0.829 | 0.962 | N/D | N/D | 0.831 | 0.886 | 0.922 | 0.974 | 0.946 |
| **Refinement** | | | | | | | | | | | |
| Resolution (Å) | 50-3.0 | 44.3-2.5 | 22.1-2.9 | 46.7-3.0 | 48.9-2.5 | 48.1-2.75 | 37.9-2.95 | 48.6-2.7 | 36.4-2.7 | 20.2-2.7 | 24.5-2.9 |
| $R_{work}/R_{free}$ | 0.233/0.306 | 0.205/0.26 | 0.236/0.306 | 0.195/0.250 | 0.194/0.244 | 0.229/0.293 | 0.217/0.279 | 0.219/0.299 | 0.179/0.236 | 0.221/0.281 | 0.226/0.289 |
| RMSD | | | | | | | | | | | |
| Bond length (Å) | 0.0098 | 0.0084 | 0.0137 | 0.0098 | 0.008 | 0.0113 | 0.011 | 0.008 | 0.0095 | 0.009 | 0.010 |
| Angle (°) | 1.162 | 1.108 | 1.905 | 1.181 | 0.999 | 1.579 | 1.488 | 1.069 | 1.362 | 1.053 | 1.117 |
| Ramachandran statistics | | | | | | | | | | | |
| Favored regions (%) | 93.17 | 97.30 | 92.34 | 96.50 | 97.41 | 94.85 | 96.87 | 92.06 | 95.95 | 94.88 | 95.22 |
| Allowed regions (%) | 5.91 | 2.45 | 6.99 | 3.44 | 2.59 | 5.15 | 2.81 | 7.57 | 3.68 | 4.75 | 3.86 |
| Outliers (%) | 0.43 | 0.25 | 0.67 | 0.06 | 0 | 0 | 0.31 | 0.37 | 0.37 | 0.18 | 0.92 |
| PDB ID | 6UJG | 6UJM | 6UK6 | 6UKB | 6UL9 | 6UMC | 6ULA | 6UMF | 6ULJ | 6UMD | 6UME |

**Table S1:** Diffraction data collection and structure refinement statistics of the GAC-inhibitor complexes. Data in parentheses refer to the highest resolution shell.

| Inhibitor | BPTES | UPGL-4 |
|---|---|---|
| **Data collection** | | |
| Space group | P21 | P21 |
| Cell dimensions | | |
| a (Å) | 50.0 | 53.2 |
| b (Å) | 138.00 | 138.44 |
| c (Å) | 178.00 | 177.49 |
| β (°) | 90.00 | 93.44 |
| Resolution (Å) | 20-3.0 | 20-2.8 |
| Unique reflections | 47136 | 61831 |
| | (4690) | (6026) |
| Redundancy | 19.8 (19.8) | 27.4 (22.2) |
| Completeness (%) | 97.6 (98.6) | 97.9 (97.1) |
| Average I/σI | 5.4 (1.3) | 5.0 (0.5) |
| $R_{merge}$ | 0.468 (2.34) | 0.670 (23.23) |
| $R_{meas}$ | 0.496(2.39) | 0.678 (23.80) |
| CC1/2 | 0.970 | 0.972 |
| **Refinement** | | |
| Resolution (Å) | 20-3.0 | 20-2.8 |
| $R_{work}/R_{free}$ | 0.207/0.288 | 0.209/0.278 |
| RMSD | | |
| Bond length (Å) | 0.009 | 0.010 |
| Angle (°) | 1.189 | 1.185 |
| Ramachandran statistics | | |
| Favored regions (%) | 91.84 | 92.06 |
| Allowed regions (%) | 6.71 | 6.75 |
| Outliers (%) | 1.44 | 1.19 |
| PDB ID | 7RGG | 7REN |

**Table S2**: Serial diffraction data and structure refinement statistics of the GAC-inhibitor complexes. Data in parentheses refer to the highest resolution shell.

| Descriptors for KPLS Model | Descriptor Meaning |
|---|---|
| **Del(K)Min** | **Rate of change in K electronic kinetic energy normal to molecular surface** – minimum value of Del(K) over entire molecule |
| Del(K)NA1 | Histogram bin 1, total surface area with low values of Del(K) |
| Del(K)NA3 | Histogram bin 3, total surface area with intermediate-low values of Del(K) |
| Del(K)NA8 | Histogram bin 8, total surface area with high values of Del(K) |
| SIKMin | **Surface Integral of Del(K)** – minimum value over entire molecule |
| SIKA3 | Histogram bin 3, total surface area with intermediate-low values of SIK |
| SIKA8 | Histogram bin 8, total surface area with high values of SIK |
| Del(G)NMax | **Rate of change in G electronic kinetic energy normal to molecular surface –** Maximum value over entire molecule |
| **Del(G)NA2** | Histogram bin 2, total surface area with low values of Del(G) |
| **Del(G)NA4** | Histogram bin 4, total surface area with intermediate-low values of Del(G) |
| **SIEPMax** | **Surface integral of electrostatic potential –** Maximum value over entire molecule |
| **SIEPA4** | Histogram bin 4, total surface area with intermediate-low values of SIEP |
| SIEPA5 | Histogram bin 5, total surface area with intermediate values of SIEP |
| EP7 | **Electrostatic Potential** – Histogram bin 7, total surface area with intermediate-high values of EP |
| **PIP4** | **Politzer's average local ionization potential –** Histogram bin 4, total surface area with intermediate-low values of PIP |
| FukMin | **Fukui radical reactivity index** – Minimum value over entire molecule |
| **LaplMax** | **Laplacian of the electron density** – Maximum value over entire molecule |
| FDRNA6 | **Fractional Del(Rho)NA6 –Rate of change of Rho kinetic energy normal to molecular surface, divided by total surface area (size invariant)** – Histogram bin 6, fractional total surface area with moderate values of Del(Rho)N |
| FDRNA10 | Histogram bin 10, fractional total surface area with high values of Del(Rho)N |
| FDKNA5 | **Fractional Del(K)NA6**, Histogram bin 5 |
| **FDKNA8** | Histogram bin 8 |
| FSIKA3 | **Fractional SIK**, Histogram bin 2 |
| FSIGA8 | **Fractional SIG, Surface Integral of G kinetic energy density**, Histogram bin 8 |
| **FEP3** | **Fractional EP/Electrostatic Potential**, Histogram bin 3 |
| **FEP5** | Histogram bin 5 |
| FEP6 | Histogram bin 6 |
| FPIP3 | **Fractional PIP**, Histogram bin 3 |
| **FPIP4** | Histogram bin 4 |
| Ffuk9 | **Fractional Fuk**, Histogram bin 9 |

**Table S3**: TAE descriptors used for KPLS model of BPTES-class inhibitory potency. In general, descriptors describe either 1) the minimum or maximum value for a given electronic calculation over a molecular surface (labeled with Min or Max at end), 2) the total surface area of the molecule having a value for a given descriptor inside a certain range (labeled 1-10 for each of 10 histogram bins), or 3) the relative amount of surface area for a molecule with a certain range of values (labeled with an initial F for 'fractional', and ending with a 1-10 to designate histogram bin). Descriptors which were positively correlated with activity in the model are written in bold green letters. A more robust explanation of each descriptor, including the formulae used to calculate them, is available in the literature (22, 28).

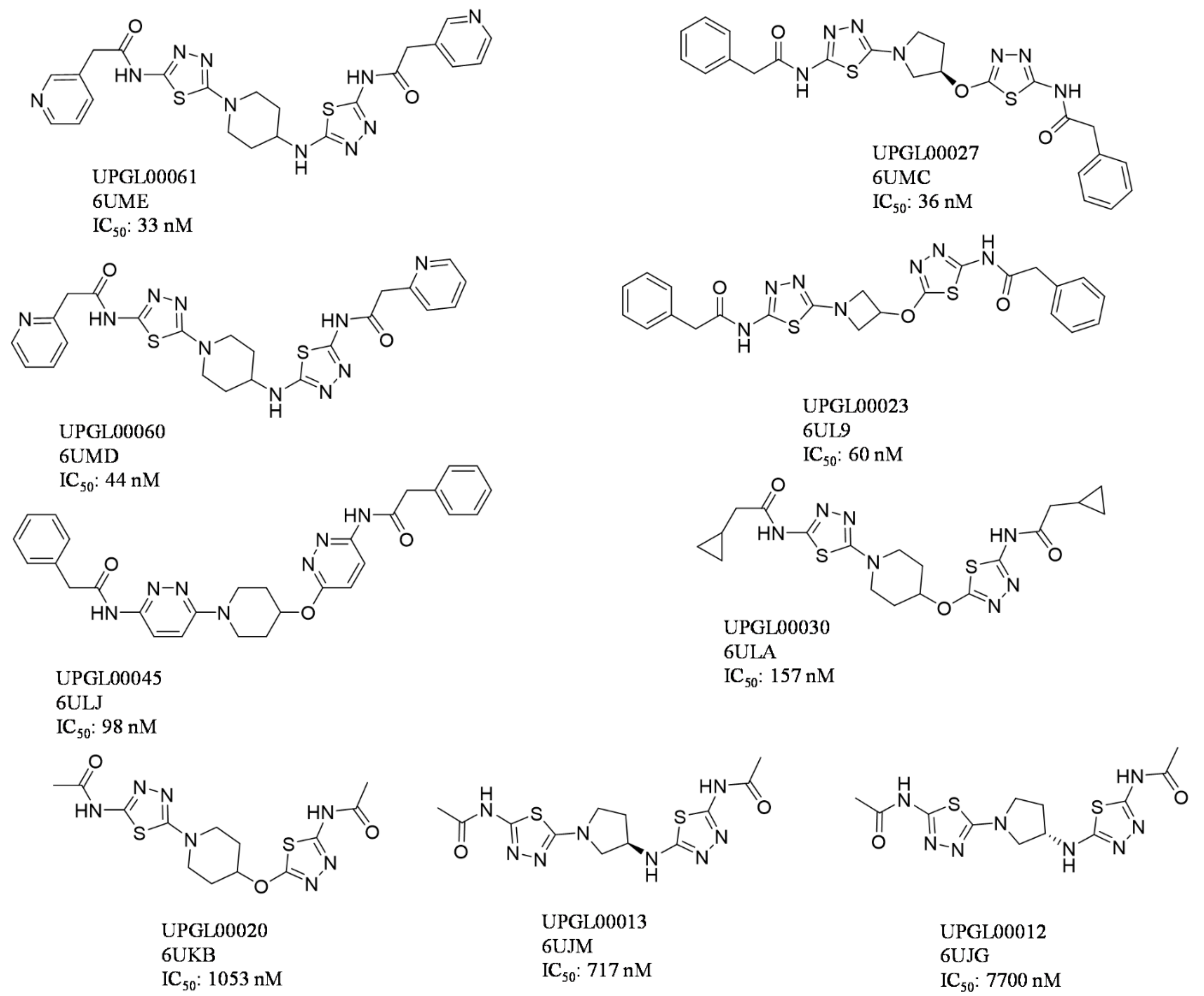


**Figure S1:** UPGL series inhibitors co-crystalized with GAC in this study. For each compound, its PDB ID for co-crystal structure with GAC, and IC50 as determined in reference (1) are reported.

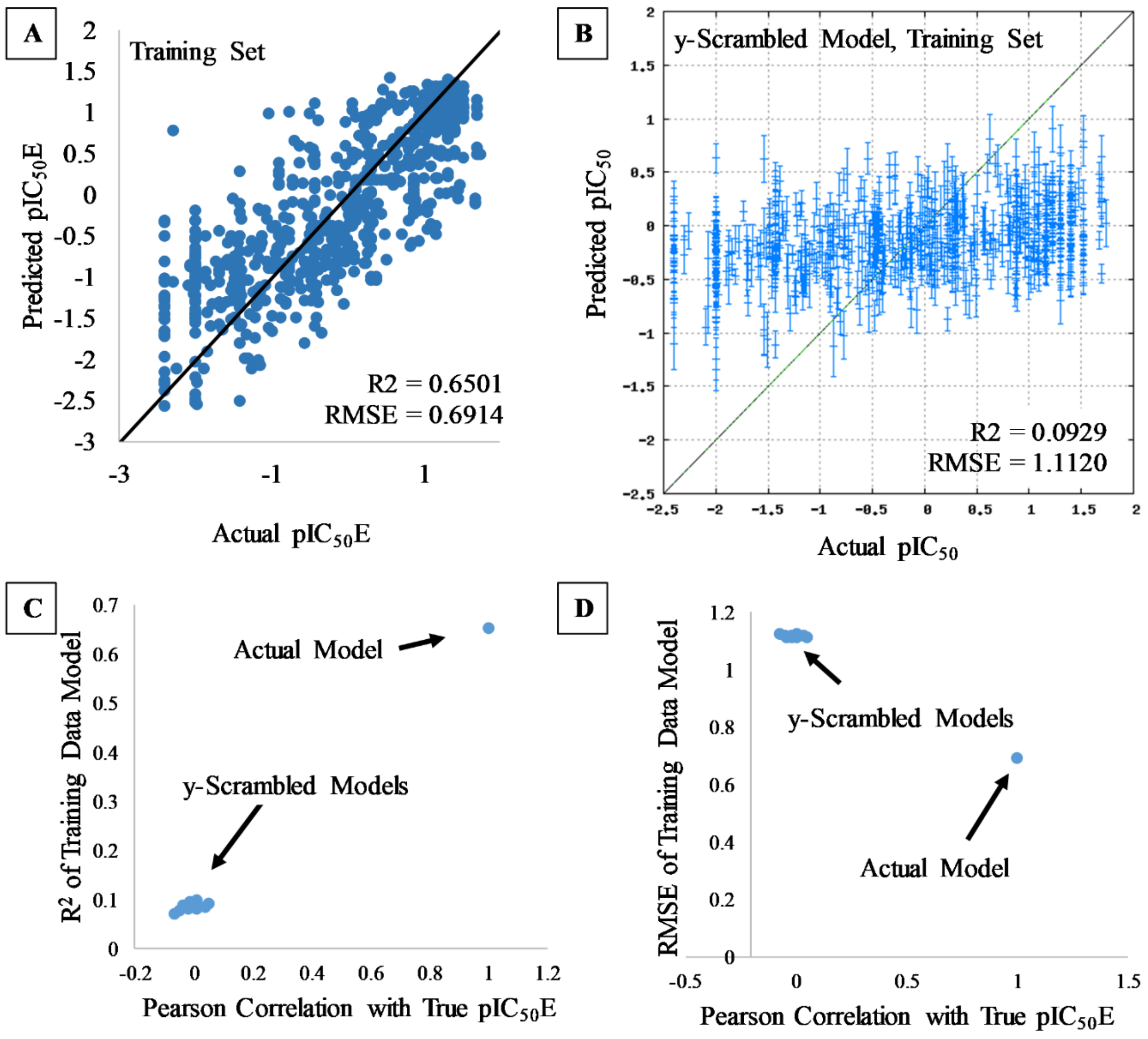


**Fig. S2**: Additional QSAR modeling of the BPTES/CB-839-class inhibitors. A) KPLS model training set results. These are the training data for the model shown in Fig. 5A in the main text. B) Y-scrambled training data, modeled by KPLS using the same configuration as in (A). The model is entirely non-predictive. C) $R^2$ of 10 y-scrambled KPLS models vs. the correlation between the scrambled $pIC_{50}E$ values and the true values. A data point is also shown for the KPLS model built with unscrambled data. D) RMSE vs. correlation for the same models shown in (C).

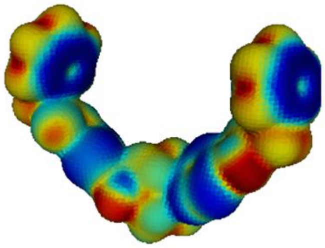

UPGL00004, $IC_{50}$ = 29 nM

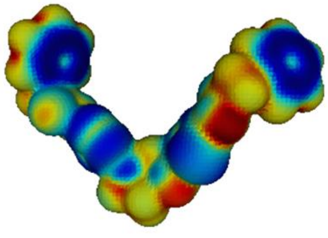

UPGL00011, $IC_{50}$ = 30 nM

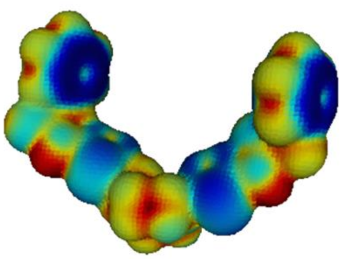

UPGL00019, $IC_{50}$ = 30 nM

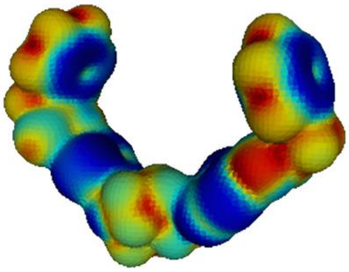

UPGL00061, $IC_{50}$ = 33 nM

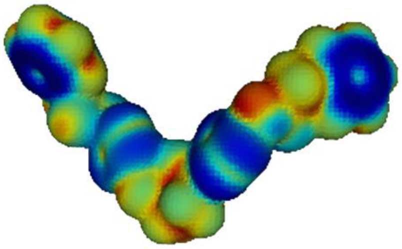

UPGL00027, $IC_{50}$ = 36 nM

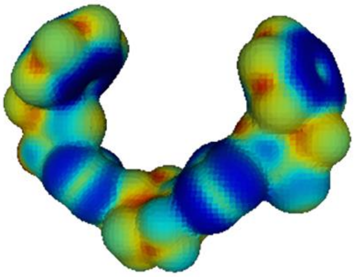

UPGL00015, $IC_{50}$ = 40 nM

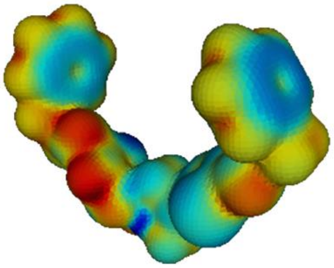

UPGL00023, $IC_{50}$ = 60 nM

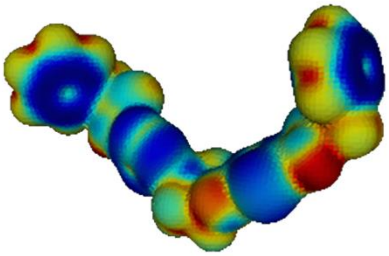

UPGL00009, $IC_{50}$ = 70 nM

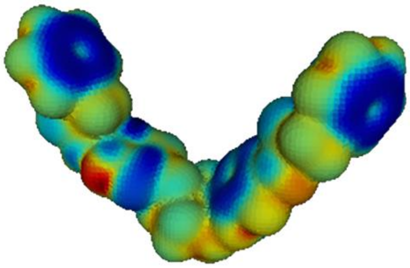

UPGL00045, $IC_{50}$ = 98 nM

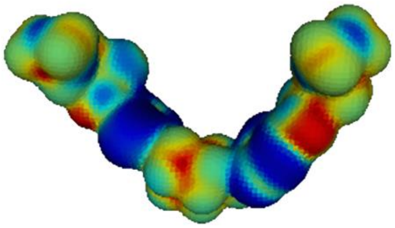

UPGL00030, $IC_{50}$ = 157 nM

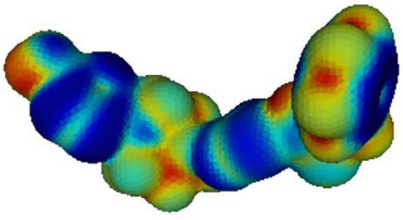

UPGL00031, $IC_{50}$ = 203 nM

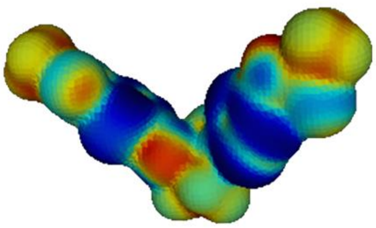

UPGL00013, $IC_{50}$ = 717 nM

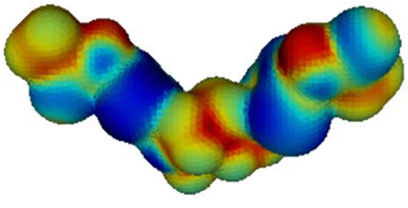

UPGL00020, $IC_{50}$ = 1053 nM

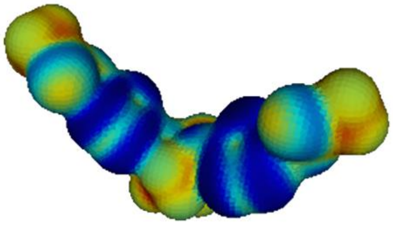

UPGL00012, $IC_{50}$ = 7700 nM

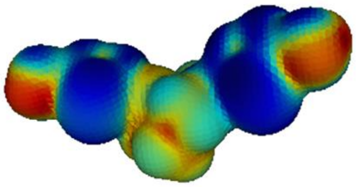

UPGL00018, $IC_{50}$ > 10000 nM

**Figure S3:** Politzer's average local ionization potential surface maps for UPGL compounds which have been co-crystallized with GAC. Compounds were extracted from the relevant PDB files for calculations. Blue regions are more easily ionizable, while red regions resist ionization. Coloration is internally scaled for each molecule. Molecules are organized by potency as measured against recombinant GAC, taken from (1).

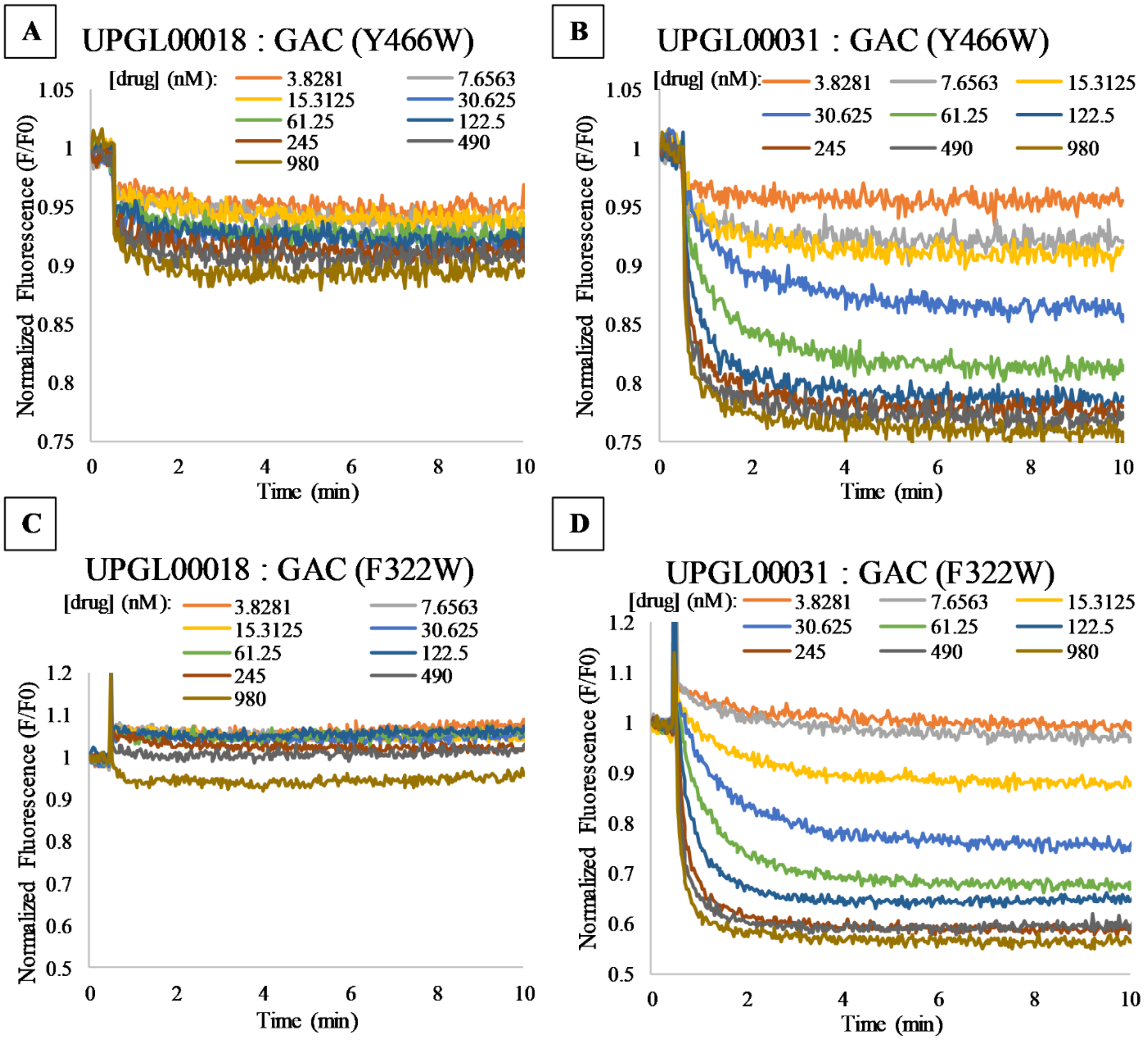


**Figure S4**: Raw data from fluorescent binding assays. A) Real-time tryptophan fluorescence emission signal ($\lambda_{ex}$ = 285nm, $\lambda_{em}$ = 340nm) of 100 nM GAC (F322W) is quenched upon the addition of (A) UPGL00018 or (B) UPGL00031, and the fluorescence signal of 100 nM

GAC(Y466W) treated with (C) UPGL00018 or (D) UPGL00031 are shown. Data is representative of three independent experiments.
